# Hepatic isomiR landscaping reveals new biological insights into metabolic dysfunction in steatotic liver disease

**DOI:** 10.1101/2025.04.08.647685

**Authors:** Christian Brion, Stephen A. Hoang, Guangliang Wang, Faridodin Mirshahi, Jessie Ang, Matthew R. Long, Zheng Zhu, Bhanu Sakhamuri, Molly A. Srour, Mohammad S. Siddiqui, Amon Asgharpour, David J. Hayes, Neal C. Foster, David W. Salzman, Arun J. Sanyal

**Affiliations:** Gatehouse Bio Inc.; 22 Strathmore Road, Natick, MA, 01760, USA; Division of Gastroenterology, Hepatology and Nutrition, Department of Internal Medicine, Virginia Commonwealth University School of Medicine; Richmond, VA, 23298, USA

**Keywords:** MicroRNA, IsomiRs, MASLD, Fibrosis, Liver Disease

## Abstract

Post-transcriptionally modified microRNA (miRNA), called isomiRs, expand the repertoire of transcripts that can leveraged for therapeutic targets and biological insights. However, the expression of isomiRs has not been characterized in metabolic dysfunction-associated steatotic liver disease (MASLD). Therefore, we assessed the isomiR expression profile in liver biopsies from 79 patients with MASLD and modeled their potential role in disease biology. MiRNAs represented 75% of the sequencing reads and over 65% of them were attributed to isomiRs, demonstrating their higher expression and diversity compared to canonically annotated miRNAs. Differential expression and machine-learning analyses were used to identify 173 isomiRs associated to MASLD severity and 58 isomiRs associated to fibrosis score. Candidate target mRNAs were identified for each isomiR based on sequence complementarity. Using matched mRNA sequencing data, and supported by data from an independent study, we proposed key dysregulated mRNA targets involved in a selection of 33 disease-associated pathways. Importantly, isomiRs offered novel and unique mRNA targets compared to the canonical miRNA, e.g. isomiR-122 targeting INSIG1 (insulin and cholesterol metabolism), and isomiR-21 targeting HMGCS2 and PPARA (PARR and TGF-beta signaling). Our work advances knowledge regarding the role of isomiRs in MASLD and lays a foundation for therapeutic targets identification.

**Highlights:** Our results provide a comprehensive analysis of microRNA (isomiRs) in liver tissue. Machine learning identified isomiRs whose expression is associated with MASLD. Multi-omic analysis uncovered novel isomiR regulatory mechanisms involved in MASLD.

## INTRODUCTION

Metabolic dysfunction associated steatotic liver disease (MASLD) is a leading cause of liver-related morbidity and mortality and also increases the risk of cardiac and cancer related deaths thus contributing to the overall burden of non-communicable diseases globally (1–3). It two principal histological phenotypes including a metabolic dysfunction associated steatotic liver (MASL) and metabolic dysfunction associated steatohepatitis (MASH). MASH is a more active form of the disease and progresses to cirrhosis and end-stage liver disease more readily than MASL.

MASLD is a heterogeneous disorder resulting from complex gene-environment interactions. The heterogeneity is likely responsible for variable clinical and histological response to therapy despite increasingly more effective drugs such as GLP-1 receptor and FGF-21 receptor agonists and the recent approval of resmetirom a thyroxine beta receptor agonist for MASH (4–7). Furthermore, some individuals do not respond to therapy while others are intolerant of therapy. These underscore the continued need to better understand the causes of disease heterogeneity to strengthen the ability to develop Precision approaches for therapeutics.

A source of heterogeneity is the variable gene expression patterns across individuals who have otherwise relatively indistinguishable histological severity and pattern of disease (8, 9). The translation of gene expression to proteins connects genetic and transcriptomic drivers of disease to their downstream effects on metabolism, inflammation, and fibrosis the key pathophysiological aspects of MASLD (9–11). MicroRNAs (miRNAs) modulate this process with multiple miRNAs targeting a given gene and multiple genes being affected by a single miRNA (12). We originally described the miRNA profile of MASLD, which used to be called nonalcoholic fatty liver disease (NAFLD), almost two decades ago (13). Since then, multiple miRNAs have been linked to liver disease through their activity in lipid metabolism (e.g. miR-122 and miR-34a), inflammation (e.g. miR-22, miR-122), fibrosis (e.g. miR-10b-5p, miR-132, miR-21) and oncogenesis (e.g. miR-22, miR-221, miR-122, miR-132) (14–23). The two most widely documented miRNAs involved in liver disease are liver-specific miR-122, suggested to regulate cholesterol and fatty acid synthesis (15, 16) and proposed as an important potential blood biomarker for liver damage (17), and miR-21, regulating inflammation and fibrosis through inhibition of the peroxisome proliferator-activated receptor (PPAR) pathway leading to potential implication in liver disease (18, 19) and ischemia (21, 24). However, therapeutic investigations of other miRNAs led to clinical trials such as those involving antago-miR-22-5p (Resalis Therapeutics) and anti-miR-132-3p (Regulus Therapeutics) (25, 26).

Over the last few years, there has been growing interest in miRNA alternative sequences, called isomiRs, that arise through 5’- and 3’-templated and non-templated additions and deletions (27). These isomiRs may differentially impact the function of various miRNAs (27). This raises the possibility of leveraging this to better understand the heterogeneity of miRNAs and other small RNAs (sRNA) and their role as disease drivers and leverage these to develop more precisely targeted therapeutics impacting both liver-based drivers of MASLD progression and MASLD related drivers of cardio-renal-metabolic disorders in the future. Previous studies have looked at the role of isomiRs in lipid metabolism, cardiovascular risk profile in MASLD and in cirrhosis (28–30) but did not measure associations across the histological spectrum of MASLD or disease activity. We therefore undertook this study with the goals to (i) define the spectrum of sRNAs including isomiRs across the histological spectrum of MASLD, (ii) link specific isomiRs to disease activity and fibrosis stage, and (iii) analyze the potential gene targets of these specific differentially expressed isomiRs and the potential pathways impacted by these isomiRs to explore hitherto unexplored regulatory maps underlying heterogeneous activity and fibrosis profiles in MASLD.

## MATERIALS AND METHODS

### MASLD patients panel and liver samples

Patients with MASLD attending the clinics at Virginia Commonwealth University Health System were approached to provide consent to have blood and liver tissue stored in a biorepository to support translational studies on liver diseases. Liver tissue was obtained from needle biopsy specimens from those undergoing a clinically indicated liver biopsy to evaluate their liver disease. The liver tissue obtained was first inspected visually and at least 2 cm length sent to clinical pathology for clinical evaluation. In individuals who provided informed consent to participate in the biorepository study, any remaining tissue was put in pre-labeled cryovials and snap-frozen in liquid nitrogen at the bedside within five minutes of the biopsy being performed. The flasks of liquid nitrogen were later transported to the investigators laboratory where the vials were put in a −80° C freezer. Samples were removed and transported to Gatehouse Bio for RNA isolation and sRNA/mRNA analyses under dry ice thus preventing thawing prior to its analysis. Clinical, demographic and laboratory data were obtained from clinical medical records.

### Histological Assessment of MASLD

The clinically indicated liver biopsy sections were read by a clinical liver pathologist with experience in reading liver biopsies and scoring MASLD related parameters. Four micrometer formalin-fixed paraffin-embedded sections were stained with either hematoxylin and eosin. or Masson trichrome. The presence of steatosis, hepatocellular ballooning, lobular inflammation and fibrosis were noted and scored using the NIDDK NASH CRN pathology scoring system as previously described (31, 32). The NAFLD activity score (NAS) is a composite and sum of the scores for steatosis, lobular inflammation and hepatocellular ballooning and was computed from the individual scores for these parameters (31). Fibrosis was assessed from the Masson trichrome stain and staged from 0-4, with four representing cirrhosis-level fibrosis and zero, no fibrosis (Table S1) (33).

### Total RNA extraction

A total of 10 mg of frozen liver tissue was homogenized using TissueLyser II (QIAGEN) with 5mm metal beads (QIAGEN). Total RNA was extracted from homogenized tissue using miRNeasy Tissue/Cells Advanced Kits (QIAGEN) following manufacturer’s instructions. To evaluate the quality of total RNAs, RINs were determined using the LabChip GX Touch Nucleic acid analyzer RNA assay (PerkinElmer). Quantities of total RNA were measured with the Quant-IT RNA assay kit (Thermo Fisher Scientific) and a Varioskan microplate reader (Thermo Fisher Scientific).

### Small RNA sequencing

Small RNA libraries were constructed using NEXTFlex Small RNA-Seq Kit v3 (PerkinElmer) and SciClone NGSx iQ Workstation (PerkinElmer). Across samples, RNA quantities were normalized to 300ng in 10.5 uL total volume. Size selection of 140–200 nucleotides fragments were isolated using a 3% agarose gel and a PippinHT (Sage Science). The final library quality was determined using the LabChip GX Touch with an NGS 3K DNA Assay (PerkinElmer). The quantities of the final library were measured with the Quant-IT 1x dsDNA Assay Kit (Thermo Fisher Scientific) and Varioskan microplate reader (Thermo Fisher Scientific). Libraries were pooled at various volumes aiming at a normalized final concentration of 1.6 nM per sample. The pooled library was sequenced on a NovaSeq 6000 (Illumina) following the manufacturer’s instructions.

Sequencing reads were processed by trimming adaptor sequence using an in-house Regex-based search and trim algorithm. For 3’ adaptor identification, the sequence 5’ TGGAATTCCTCGGGTGCCAAGG 3’ (containing up to a 15 nucleotide 3’-end truncation) was input; and a Levenshtein Distance of 2 or Hamming Distance of 5. Parameters for Regex searching required that the 1^st^ nucleotide of the 3’-adaptor be unaltered with respect to nucleotide insertions, deletions, and/or swaps. For 5’-adaptor identification, the sequence 5’ TCTTTCCCTACACGACGCTCTTCCGATCT 3’ (containing up to a 15 nucleotide 5’-end truncation) was input, and a Levenshtein Distance of 2 or Hamming Distance of 5. Parameters for Regex searching required that the 29^th^ nucleotide of the user-specified search term be unaltered with respect to nucleotide insertions, deletions, and/or swaps. Paired-end reads were removed if they were not an exact match. A 4-nucleotide NNNN prefix and NNNN postfix, inherent in the Nextflex adaptors, were used as a Unique Molecular Index (UMI) to qualify the quality of the sequence. Sequencing data are available on NIH under BioProject number PRJNA1242911.

### Small RNA sequence mapping and isomiR annotation

The read-count per UMI ratios were used to estimate levels of PCR duplicates, and sRNA with lower-than-expected level of PCR duplicate were flagged as potentially lower quality, which corresponded to 6,016 out of the 102,943 sRNA. Each individual trimmed sequence was regarded as a potential sRNA molecule (including isoforms) and mapped onto the human genome to identify its origins in two steps. First, the sequences were aligned to a set of 2632 canonical miRNA sequence—one per miRNA families—with 10 nucleotides templated extensions on both sides using bowtie I. Only two mismatches were allowed within the sRNA sequence, but additional mismatch were allowed on either the 3’ and 5’ extremities. This process resulted in many isomiRs having multimappings to multiple miRNA families. Each isomiRs mapping was then scored with penalties of 1 for TADs, 2 for NTAs and 5 for swaps with the lowest score utilized for the final mapping. The second mapping aligned the remaining unmapped sequences to a 17-95 nucleotide tiled array of the human genome (hG38) with up to two mismatches allowed, using RefSeq, piRbase, and miRbase annotation. In case of multimapping outside of miRNA, mappings were prioritized first toward piRNA, then tRNA, rRNA, genes, and finally other ncRNA.

The expression of each unique sRNA was measured using the count of successfully trimmed reads that match exactly its sequence, normalized for sequencing depth: reads per million (RPM). We measured miRNA family’s expression as the sum of RPM of all sRNA whose best mapping was to the family’s respective locus. For each sRNA, we identify the presence of outlier measurements using log2RPM expression values with two independent methods. First, outliers were flagged if the measurement is above the 90th quantile plus two times the 10-90th quantile range. The second method used inter-measurement distances: outliers were measurements above the maximal inter-measurement distance if they represent less than 10% of the data and if the maximal inter-measurement distance is more than half the total distance from median to maximum expression value. If outliers were observed for any of these two methods, the sRNA was excluded from downstream analysis, which corresponded to 32,456 out of the 102,943 sRNA from this dataset (Table S2).

### Messenger RNA library preparation and mRNA-sequencing

Libraries of mRNA were constructed using NextFlex Rapid Directional RNA Seq Kit (Perkin Elmer) and SciClone NGSx iQ Workstation (Perkin Elmer). A total of 300ng of RNA were used in a 10.5 µL total volume. The final library quality was determined using the LabChip GX Touch Nucleic Acid Analyzer with a NGS 3K DNA Assay (Perkin Elmer). The quantities of the final library were measured with the Quant-IT 1x dsDNA assay kit (Thermo Fisher Scientific) and Varioskan microplate reader (Thermo Fisher Scientific). Libraries were normalized and pooled at 1.6nM. Libraries were sequenced on a NovaSeq 6000 (Illumina) using an S4 flow cell.

Paired-end reads from mRNA-seq were trimmed using Trimmomatic (34) and mapped on human hg38 genome using HiSat2 (parameters: -p 5 --dta --rna-strandness) (35). Gene expression was quantified by read count using HTseq-count (parameters: -r name -f bam -m union -s reverse --nonunique none) (36).

### Differential expression analysis

Differential expression was performed on the raw read count in R using ‘DEseq2’ package (37). In all our analyses (sRNA and mRNA), we incorporated the RIN as covariate to remove potential bias from RNA quality. A Benjamini-Hochberg multiplicity correction was applied (38), and 0.05 was used as p-adjust significant threshold for our sRNA analyses and p-adjust < 0.1 for our mRNA analyses. Differential expression data are provided in Table S2 (sRNA) and S6 (mRNA).

### Ordinal regression analysis

For expression correlation to NAS and fibrosis score, an ordinal regression method was deployed in R using ‘Ordinal’ package. A cumulative link logit model as fit for each sRNA or genes, as described by Hoang et al. (9), and similarly to this study, the ordinal regression reported fold changes were calculated as the difference in the mean log2RPM between the top two and bottom two levels of either NAS or fibrosis score. Ordinal regression data are provided in Table S2 (sRNA) and S6 (mRNA).

### Machine learning models

All models were developed over a 10X k-fold cross-validation to reduce overfitting and to allow for the estimation of predictive—and thus diagnostic—performance. Within each fold, the top 10,000 sRNA features were first selected based on criteria that prioritize sRNA features with differential expression between classes. The sRNA features were then further reduced using ridge and lasso regression with the ‘glmnet’ R package using α=0.5, varying λ such that we achieve the desired number of selected features. Finally, a support vector machine (SVM) was fit to the training set using a linear kernel. Descriptive statistics including Shapley Additive Explanation (SHAP) values were calculated for these models to identify the individual feature importance. The model was then applied to the testing set and performance metrics including accuracy, positive and negative predictive values, sensitivity and specificity, and receiver operator characteristic area under the curve (AUROC) were calculated for each model. Performance metrics from the combined results of the cross validation were also calculated, with the AUROC being combined using the ‘ROCR’ package’s prediction function.

The target number of features from elastic net regression was selected by choosing the best performing models, as determined by mean AUROC, containing the fewest maximum number of unique sRNAs. Based on this approach, we selected up to 27 features per fold for NAS high vs low, 40 features for fibrosis high vs low, and 66 features for MASL vs MASH (Table S3-4). For all machine learning models, sequencing data was linearized by utilizing log2-transformed RPM as the measurement.

### Gene set enrichment analysis

Gene enrichment analysis for both differentially expressed genes and TargetScan target list were done in R using ‘ClusterProfiler’ and ‘ReactomePA’ (39, 40) on Biological Process from Gene Ontology, Kyoto Encyclopedia of Genes and Genomes (KEGG) pathway, and Reactome pathway database. Only pathways with between 3 and 300 genes were considered. Significance was defined at a threshold of 0.05 on the adjusted p-values after Benjamini-Hochberg (FDR) correction (38).

### AI-generated Knowledge Graph for miRNA-Disease-Gene-Pathway associations

To construct a dynamic knowledge graph, multiple Large Language Models (LLMs), including Claude, Perplexity, and ChatGPT, were used to analyze and extract relationships between miRNAs, diseases, genes, and pathways from scientific literature from PubMed and other publication repositories (Tables S5, S8 and S9). These models were deployed in four steps.

1. Data Extraction and Processing

The AI-driven system systematically parsed through scientific publications, extracting relevant information on miRNAs, diseases, genes, and pathways. This process involves: (i) Literature Mining: The LLMs analyze abstracts and full-text articles from PubMed and other repositories, focusing on NASH-related miRNA studies. (ii) Entity Recognition: The system identifies, and extracts mentions of miRNAs, diseases, genes, and pathways using advanced natural language processing techniques. (iii) Relationship Extraction: The LLMs infer and extract relationships between the identified entities based on the context and content of the publications.

1. Knowledge Graph Construction

The extracted information is used to construct a dynamic knowledge graph with each unique miRNA, disease, gene, and pathway represented as a node and relationships between them established as edges.This method offers a knowledge graph continuously updated as new publications are processed, ensuring that the most current information is incorporated.

1. API Development

The constructed knowledge graph is exposed through an API that provides three main association lists: miRNA-to-Disease Association, Disease-to-Genes Association, and Disease-to-Pathways Association. In the context of this study, “liver” and “steatohepatitis” were used as disease key words for identifying association of miRNA to liver disease and “fibrosis” and “cirrhosis” for association to fibrosis. For both genes and pathways, we used “NAFLD” and “liver fibrosis” for association to MASH severity and advanced fibrosis, respectively.

1. Validation and Quality Control

To ensure the accuracy and reliability of the knowledge graph, results are compared automatically across multiple LLMs to identify consistencies and resolve discrepancies. Additionally, periodic manual reviews are conducted to validate the extracted relationships and overall graph structure. The system’s performance is evaluated using standard metrics such as precision, recall, and F1-score for relationship extraction tasks.

### Data visualization and additional statistical analysis

Data and statistical analysis were conducted through R language scripts (R version 4.1.3, 2022-03-10). Significance test and threshold are provided in figures and results sections.

## RESULTS

### MicroRNA is the most prominent sRNA in liver biopsies

We analyzed 79 liver biopsy samples obtained from patients with MASLD. This panel, consisting of 19 MASL, 47 MASH, 11 MASLD related cirrhosis, and 2 with MASLD but not well characterized histological pattern, exhibited varying degree of NAFLD activity score (NAS) and fibrosis stages (Fig. 1A-D, Table S1). However, since NAS declines with progression to cirrhosis (41, 42), its values for patients with cirrhosis were excluded from further analyses (Fig. 1C).

**Fig. 1.**
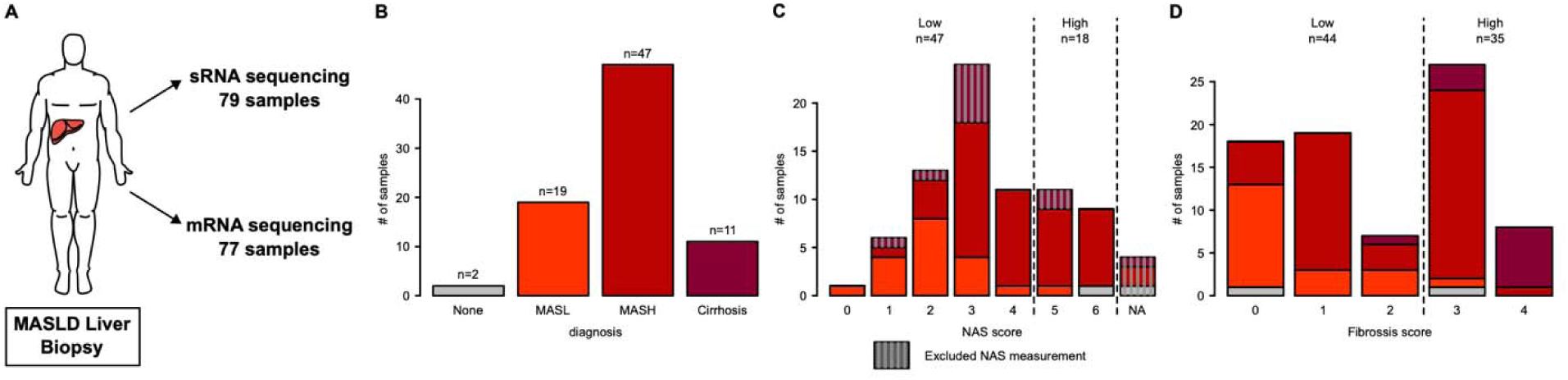
Data overview. (**A**) Samples origin and type of sequencing data. (**B-D**) Distribution of the number of samples for each key clinical information: Diagnosis (B), NAS score (Ballooning + Steatosis + Lobular Inflammation) (C), and Fibrosis score (D). The colors indicate the type of MASLD diagnosis. NAS scores for Cirrhosis patients were excluded from analysis.

Small RNA (sRNA) was extracted and sequenced from the samples, with an average 14% of the reads falling in the 17-26 nucleotides range, categorized as potential regulatory elements (Fig. S1A). Principal component analysis (PCA) based on normalized expression value (log2RPM: log2 of reads per million) showed a limited impact of diagnosis (MASL vs MASH vs cirrhosis) on the overall data structure (Fig. S1B). However, pairwise correlation analysis of the sample metadata and principal components showed a high impact of the RNA Integrity Number (RIN) on the sequencing data (Fig. S2). Other patient covariates, such as age, sex, and BMI, exhibited minimal impact on the data (Fig. S2).

Sequencing identified 102,943 unique sRNA transcripts categorized into seven distinct classes: miRNA, piRNA, tRNA, rRNA, ncRNA, and intron/intergenic regions (Table S2). The majority of sRNA transcripts mapped to miRNA (74.6%) or piRNA loci (11.3%), with sequence length primarily between 20-24 nucleotides (Fig. 2A). We sub-categorized the sRNAs originating from miRNA into their respective miRNA families. A total of 758 families were identified in this liver dataset, and among these, the 50 most highly expressed families accounted for approximately 90% of the total miRNA expression. As expected, the hepatocyte-specific miR-122-5p family was the most abundant in our dataset (41.2% of the miRNA expression), highly exceeding the second most prevalent miRNA family, miR-143-3p (5.6%, Fig. 2B and Fig. S3).

**Fig. 2.**
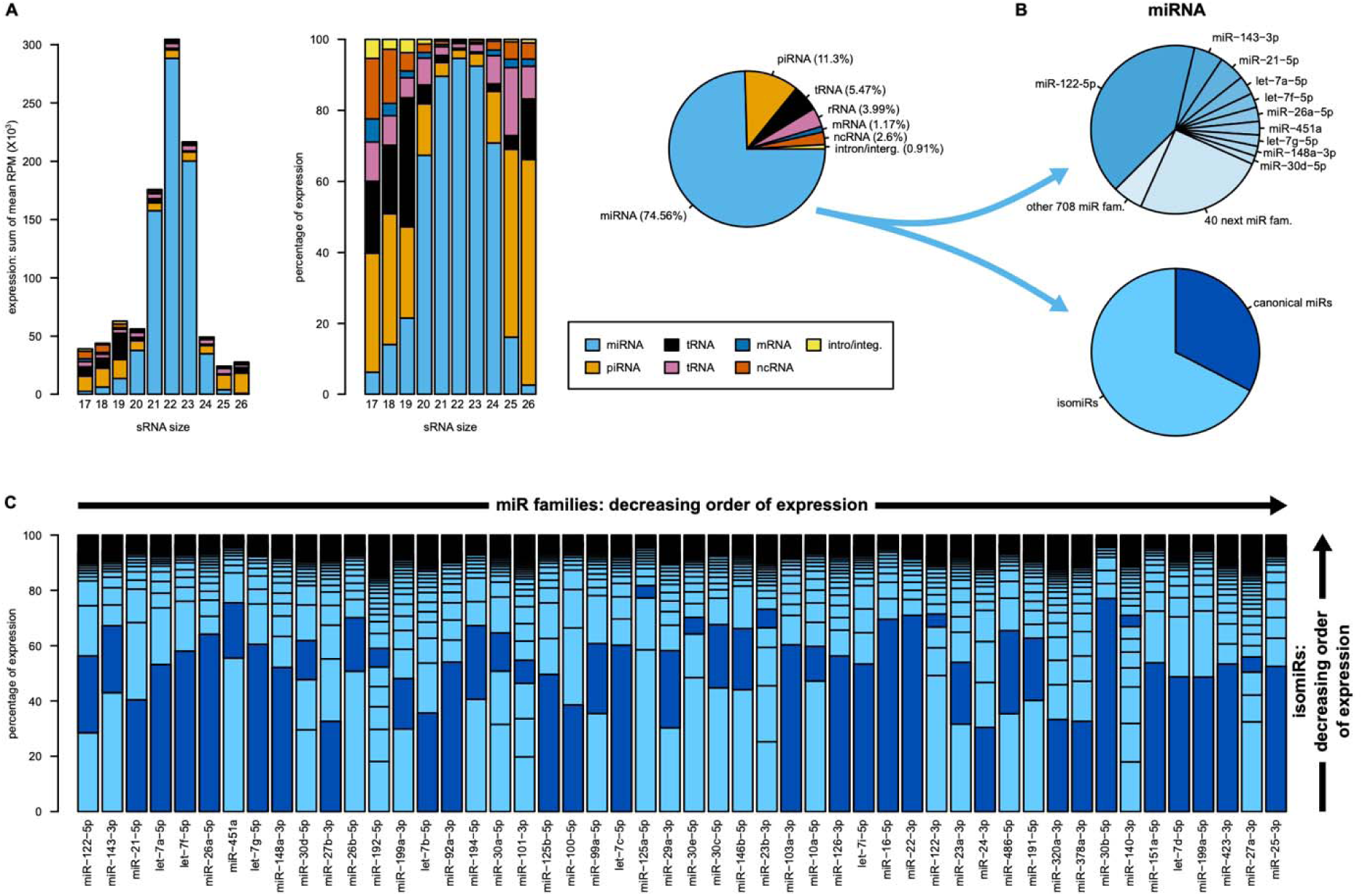
sRNA expression in liver biopsy samples. (**A**) Distribution of sRNAs size and class across samples. (**B**) Proportion of miRNA expression by families (top) and by miRNA sequence type (bottom). (**C**) The expression breakdown of the isomiRs and canonical miRNAs within the 50 most highly expressed miRNA families.

One third of the miRNA expression was attributed to the canonical sequences perfectly matching their respective miRNA family reference (Fig. 2B). The remainder, classified as isomiRs, showed potential post-translational modifications. Importantly, the number of isomiRs and their relative abundance varied widely across families. For instance, families such as miR-192-5p, miR-125a-5p, and miR-30e-5p expressed high proportions of isomiRs, where the canonical miRNA was less than 7% of the total for each family (Fig. 2C).

### MicroRNA isoforms have a significant impact on the landscape of sRNA

MicroRNA isoforms originate from alternative RNase III cleavage by DICER or DROSHA, non-templated nucleotide additions, and swaps. We classified the isomiR characteristics for each miRNA family among five types: Templated Addition or Deletion on the 5’ ends (5’ TAD) or on the 3’ ends (3’ TAD), Non-Templated Addition on the 5’ ends (5’ NTA) or on the 3’ ends (3’ NTA), and internal nucleotide swaps (Fig. 3A). For the two most abundant miRNA families, miR-122-5p and miR-143-3p, the canonical sequence is the second most expressed miRNA in the family, with the most expressed sequence carrying one 3’ NTA (Adenosine (A) addition for miR-122-5p, Uridine (U) addition for miR-143-3p, Fig. 3B). From these data, we obtained the frequency and size of these isomiR modifications. This is illustrated for these two miRNA families (Fig. 3C), highlighting the importance of the 3’ NTAs as well as 3’ TADs. The 5’ end is more conserved and less than 0.3% of transcripts had swaps between nucleotides 2-8, also referred to as the seed region (12). The same observation was made across the 50 most expressed miRNA families in our liver dataset (Fig. 3D).

**Fig. 3.**
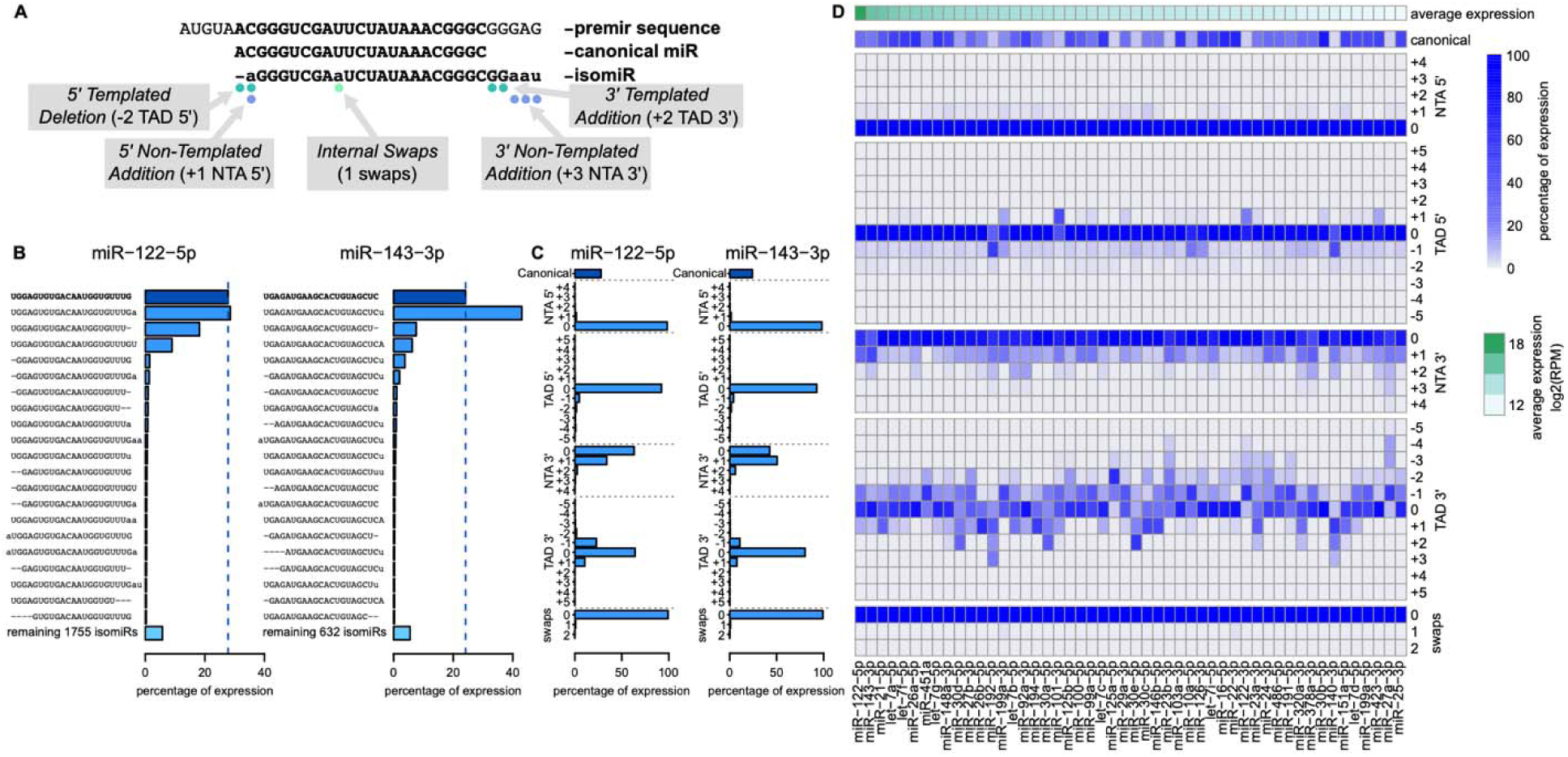
MicroRNA isoform landscape. (**A**) Overview of the five types of isomiR modifications. (**B**) Expression levels of the 20 most abundant isomiRs within the two most highly expressed miRNA families (miR-122-5p, left; miR-143-3p, right). (**C**) Distribution of the occurrence of each type of isomiR modifications within the two most highly expressed miRNA families (miR-122-5p, left; miR-143-3p, right). (**D**) Summary of isomiR modifications occurrence across the 50 most highly expressed miRNA families.

The high conservation of the 5’ end implies that changes in the seed sequence of the isomiRs compared to the canonical occur at low frequency (12% across all miRNAs). However, such alternative seeds are expected to impact the set of mRNA targets of the isomiRs and open potentially new regulatory mechanisms that could be significant in disease progression, even if the relative frequencies of these isomiRs are low. Moreover, there are some noteworthy exceptions with miR-192-5p, miR-199a-3p, miR-101-3p, miR-10a-5p, miR-126-3p, miR-122-3p and miR-140-3p having higher proportion of 5’ TADs than average (Fig. 3D). Across the 50 most expressed miRNA families, frequency of 5’ NTAs remained between 0.5 and 5% of the family expression and in most cases corresponded to a single adenosine addition (Fig. S4A). These additions have significant implications as they can influence binding to the Argonaute proteins (43) along with the potential change of the isomiR seed.

The 3’ ends of isomiRs were not as well defined, with many cases where the most frequent templated last nucleotide did not match the canonical sequence (e.g. miR-125a-5p, miR-30e-5p, miR-122-3p, Fig. 3D). While in most cases the seed region is what has been reported to trigger gene silencing, recent studies now suggest that the 3’ end has a stabilizing effect on miRNA to mRNA binding, increasing repression efficiency (44–46). Moreover, two additional functions have been proposed for the 3’ terminal nucleotide of isomiRs: dictating the cellular localization of the isomiR (47, 48) and having a stabilizing/destabilizing effect with 3’ uridines potentially triggering degradation while 3’ adenosines prevent it (49). The frequency of 3’ NTAs is on average 18.6%, much higher than of 5’ NTAs, and went as high as 57.4% for miR-143-3p isomiRs. The most frequently observed NTAs were adenosines and uridines and the proportion of each varied across the 50 top expressed families, implying different isomiR stability and localization (Fig. S4B).

### Diversity of microRNA across the histological spectrum of MASLD

By characterizing the isoform landscape of microRNA in MASLD biopsy, we wanted to highlight the potential for unexplored regulatory networks in liver tissue and used it as a basis to identify those associated with either MASH severity or liver fibrosis. Our primary goal was to identify sRNAs that are differentially expressed at increasing activity and fibrosis stages of disease. To do this, we applied two complementary approaches: first, we used differential expression analysis across three comparator groups (i) MASH versus MASL diagnosis, (ii) NAS high (>4) versus low (0–4) and (iii) advanced fibrosis (NASH CRN stages 3-4) versus low fibrosis (0–2), and second, we identified sRNAs correlated with NAS and fibrosis score using ordinal regression. In both analyses, we integrated RIN as covariate to account for variability due to differences in RNA quality. For the differential expression analysis, after Benjamini-Hochberg (BH) multiplicity correction, 299, 1824, and 7818 sRNAs were found significantly differentially expressed (DE) between MASH vs. MASL, NAS high vs. low, and fibrosis high vs. low, respectively (BH p-adjust < 0.05, Fig. 4A, Table S2). The low number of DE sRNA for MASH vs. MASL might reflect the challenges of histology-based diagnosis in accurately capturing the tissue pathophysiology and the well-known challenges of histological assessment of hepatocellular ballooning, the hallmark lesion of steatohepatitis (50, 51). The ordinal regression modeling identified 99 and 961 sRNA correlated to NAS and fibrosis respectively (BH p-adjust < 0.05, Fig. 4A). The high number of sRNAs significantly associated with advanced fibrosis in both methods suggest a potential for RNA interference in the regulation of liver scarification.

**Fig. 4.**
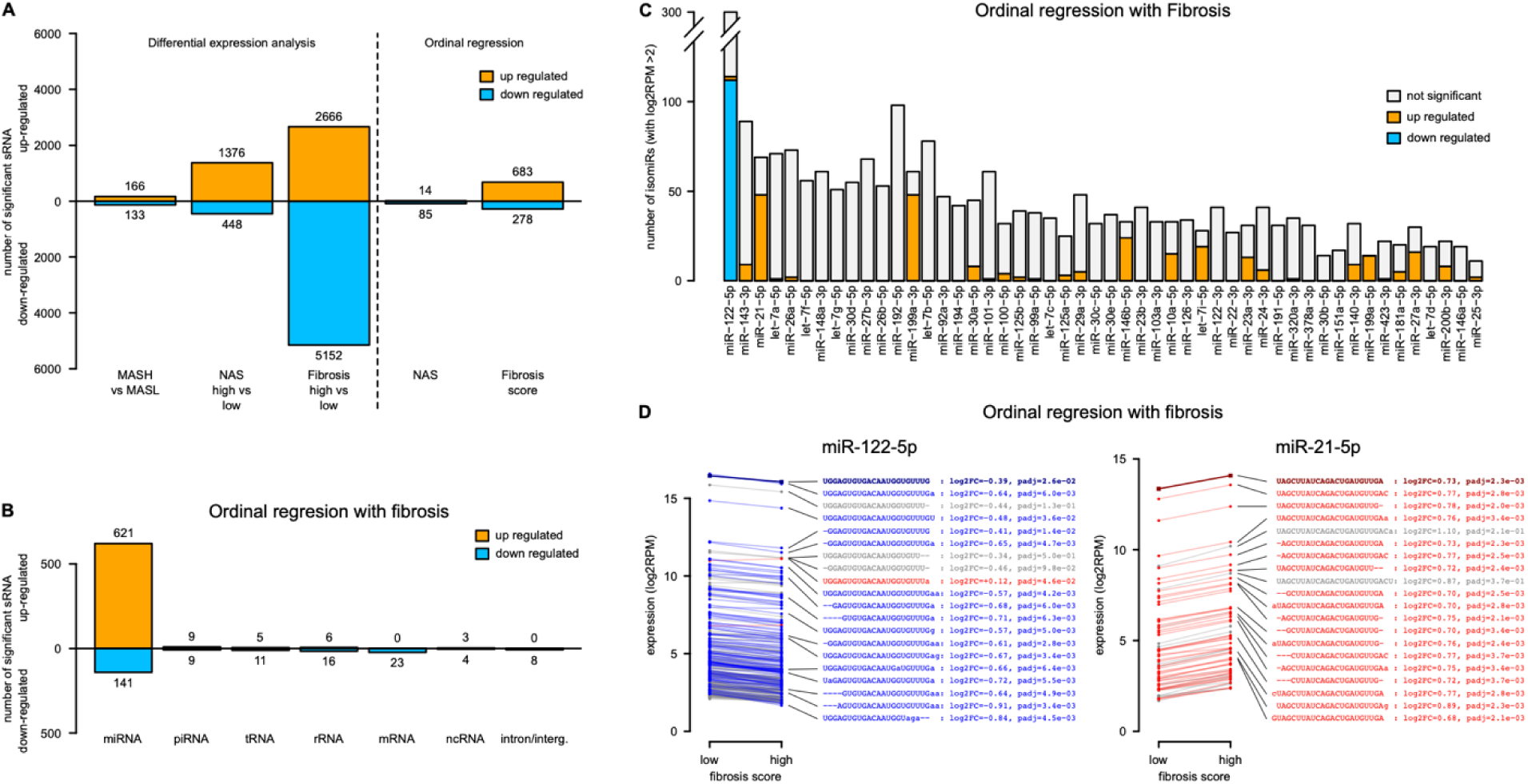
Differential expression analysis. (**A**) Number significant sRNAs identified across the three comparator groups and two ordinal regression analyses. (**B**) Breakdown of the number of significant sRNA by type from the ordinal regression analysis with fibrosis. (**C**) Number of significant isomiRs within the 50 most highly expressed miRNA families from the ordinal regression analysis with fibrosis. (**D**) Directionality and significance of all isomiRs (average log2RPM >2) within two miRNA families, left: miR-122-5p, right: miR-21-5p. The canonical miRNA sequence is highlighted in bold. Blue: down-regulated, red: up-regulated.

After filtering out sRNA with lower sequencing quality (i.e., outlier measurement or potential sequence errors) and with very low expression (i.e., log2RPM < 2.0), we looked at the distribution of the class of sRNA among the DE sRNAs (Fig. 4B and Fig. S5A) and across the 50 most highly expressed families (Fig. 4C and Fig. S5B). The liver specific miR-122-5p family contained many isomiRs negatively correlated with increasing fibrosis stage. In contrast, other dysregulated miRNA families, such as miR-143-3p, miR-21-5p, and miR-146b-5p, were positively correlated with worsening fibrosis stage (ordinal regression, Fig. 4C). We also found that most of the significantly DE miRNA in patients with high NAS scores were repressed, including isomiRs from miR-26a-5p, miR-26b-5p, miR-92a-3p and let-7c-5p (Fig. S5B). We highlighted the expression changes of miR-122-5p and miR-21-5p isomiRs, which are two highly expressed miRNA families dysregulated with advanced fibrosis and previously suggested as implicated in liver disease (15–19, 21, 24) (Fig. 4D). While most isomiRs have a consistent down- or up-regulation within the miR-122-5p and miR-21-5p families respectively, the magnitude and significance can vary. A unique case of a significantly up-regulated isomiR was observed for miR-122-5p, contrasting with the rest of the family’s isomiRs. While this sequence shares an identical 5’ end to the canonical miR-122-5p, its 3’ end was altered: one nucleotide deletion and one adenosine NTA, potentially changing the molecule stability and cellular localization. Overall, these data demonstrate substantial variability in the sRNA and isomiR landscape across the histological spectrum with substantial changes linked to advanced fibrosis.

### Machine learning highlights the involvement of isomiRs in MASLD

We further explored the relationship of liver sRNA profiles with the histological classification of MASLD patient, disease activity, and fibrosis stages by applying machine learning (ML) algorithms to our sRNA expression data. The objective was not-only to identify patterns of sRNA associated with increased disease severity and advanced fibrosis but also to gain confidence in the involvement of specific sRNA by their predictive potential through 10-fold training-testing sample split and cross-validation (Fig. 5A). For each fold, our model first down-selected sRNA features using elastic net regression and fit a linear support vector machine (SVM) to the training set which was then applied on the testing set.

**Fig. 5.**
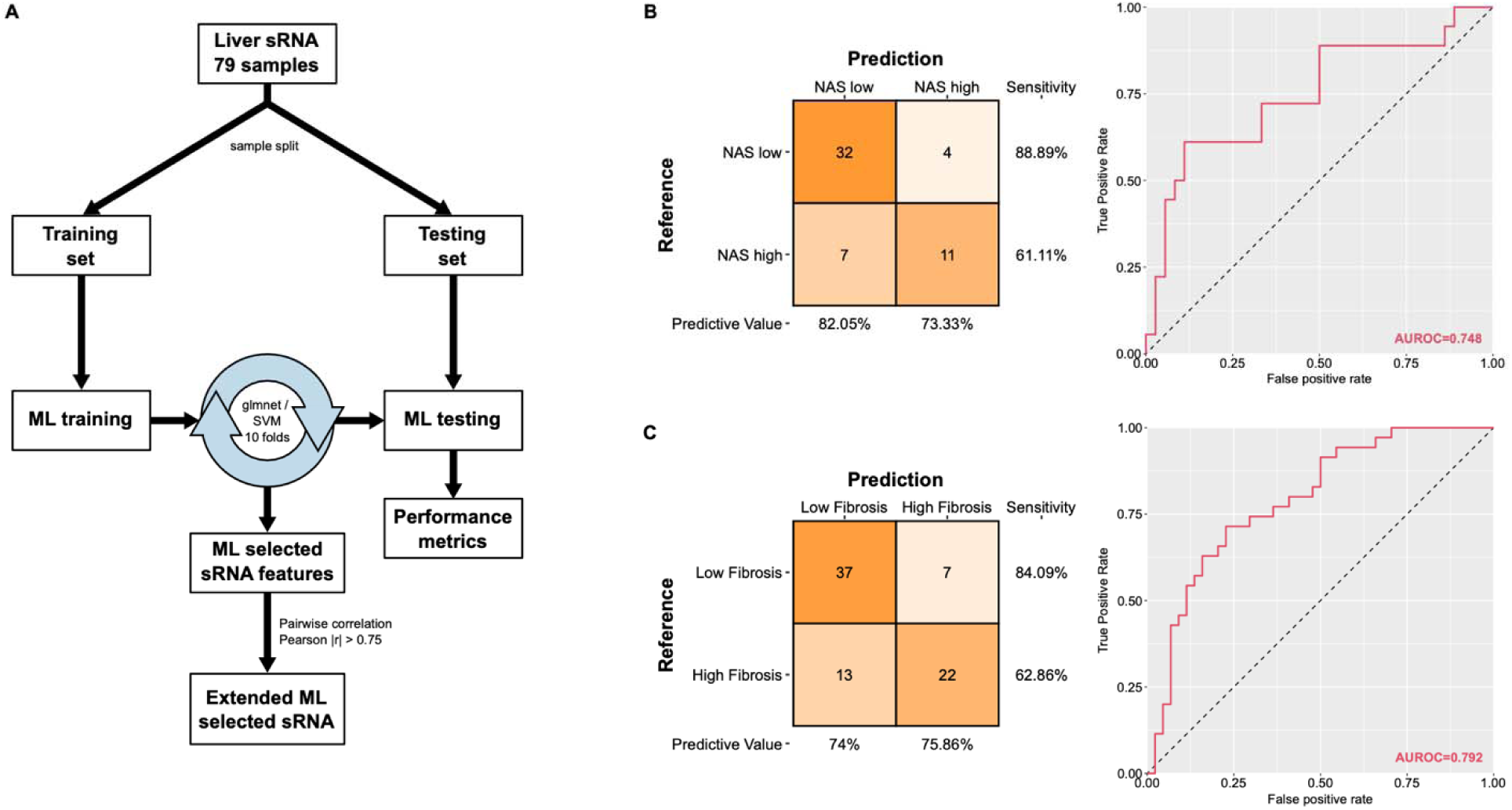
Machine Learning assisted sRNA selection. (**A**) Method overview. (**B-C**) Predictive outcomes with confusion matrix (left) and ROC curve (right) for high/low NAS scores (B) and high/low fibrosis scores (C).

The models for MASL/MASH separation did not perform well (area under the ROC curve, AUROC = 0.605, p-value = 0.303, Fig. S6), which is consistent with the limited number of DE sRNA associated with diagnosis. However, ML models for high/low NAS and Fibrosis Score had better performance (respectively AUROC = 0.748, p-value = 0.067 and AUROC = 0.792, p-value = 0.00033, Fig. 5B-C). The NAS ML models used 23 sRNA features, including isomiRs from the miR-122-5p, miR-143-3p, miR-21-5p, and miR-26a-5p families, while fibrosis models were based on only 7 sRNA features from miR-26a-5p, let-7b-5p, miR-103-3p, and miR-146b-5p (Table S3 and S4).

Since our model down-selected sRNA features, the direct ML results will only involve sRNA with distinct expression patterns and ignore sRNAs that may share similar profiles. Consequently, sRNAs with a Pearson correlation threshold of |r| > 0.75 with a feature selected by the ML model are considered as a potential predictive alternative. The resulting extended panels of transcripts were constituted of 13151 and 311 predictive sRNAs for NAS high vs low and fibrosis high vs low respectively. After filtering for sequencing quality and expression level (i.e., log2RPM > 2.0), 3948 and 268 predictive sRNAs were retained (Table S3 and S4). Finally, predictive sRNAs directly related to either MASH severity or advanced fibrosis were identified as the overlap between significant sRNAs from differential expression/ordinal regression analyses and the ML extended selection (intersections in colors on upset plots, Fig. 6A-B). For MASH and fibrosis respectively, a final list of 271 and 72 sRNAs were obtained, including 173 and 58 isomiRs across 41 and 17 miRNA families. Those families are shown in Fig. 6C-D with the number and type of isomiRs and expression directionality. Additionally, we provided for this list of miRNA families an estimate of the number of previous studies reportedly associating them with either liver disease or fibrosis using an in-house AI generated miR-family-to-disease knowledge-graph dataset (Table S5).

**Fig. 6.**
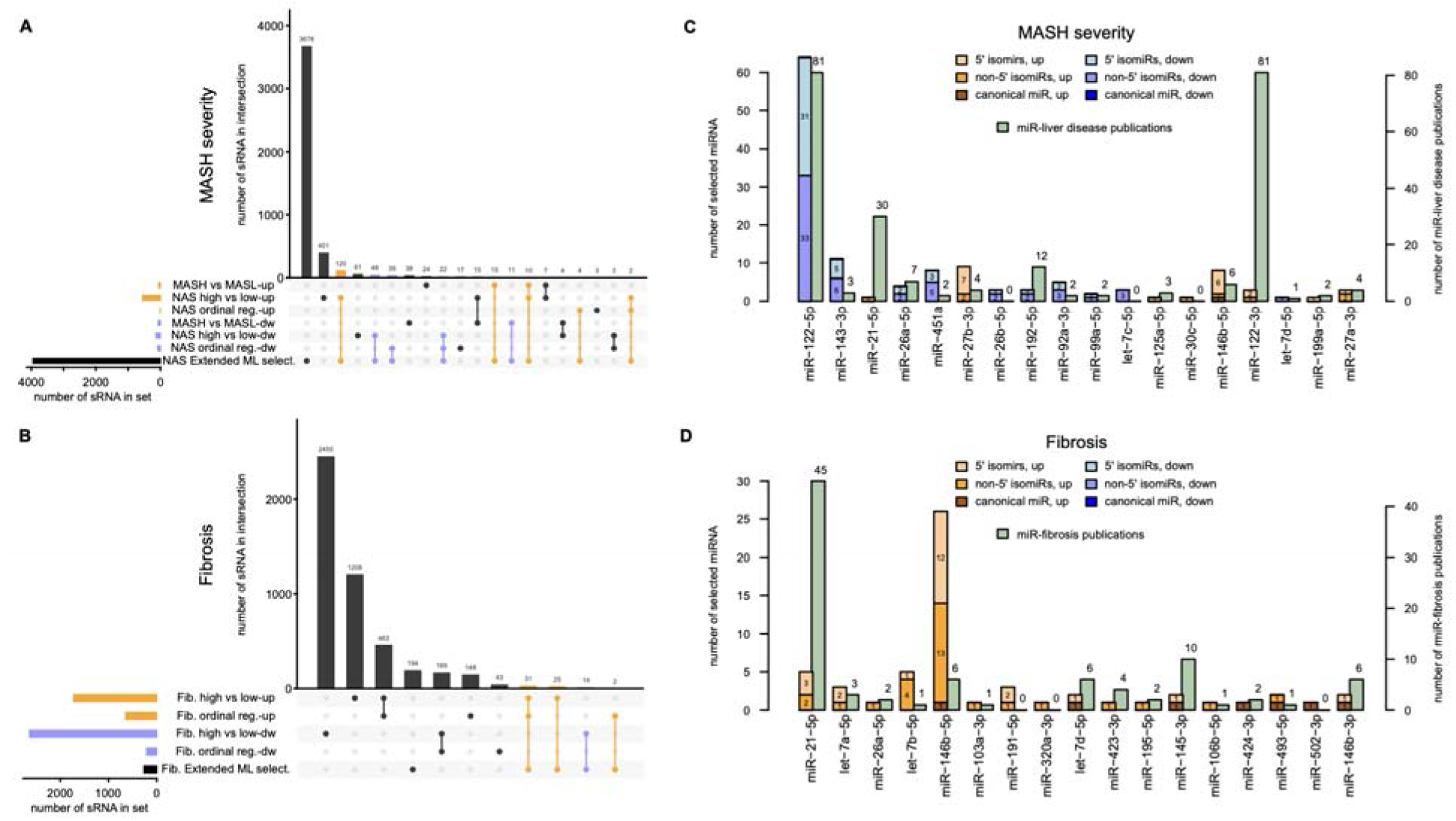
Identification of miRNA isoforms with potential disease involvement. (**A-B**) Upset plot showing differential expression analysis and machine learning (ML) outcome overlap in relation to high NAS (A) or fibrosis scores (B), across all sRNA types. (**C-D**) Summary of the isomiRs identified through both differential expression analysis and ML. The number of publications identified through an AI-generated knowledge graph associating each miRNA family to MASH severity and liver disease (C), or fibrosis (D) are also provided in the figure. MiR families are ranked left to right in decreasing order of expression. Only isomiRs from the top 50 most highly expressed miRNA families are shown in panel (C).

Importantly, our approaches not only confirmed the involvement of isomiRs from miRNA families frequently reported as associated with liver disease, including the associations of miR-122 with MASH severity and miR-21 associated with liver fibrosis (Fig. 6C-D), but also identified isomiRs from the miR-26b, let-7c, miR-30c, miR-191 and miR-320a families with no reported associations with liver disease, suggesting contribution of novel regulatory networks.

### Understanding regulatory roles of isomiRs through gene expression analysis in MASLD

To identify potential downstream regulatory gene targets and pathways that link the DE and predictive isomiRs to MASH or advanced fibrosis, we performed mRNA sequencing on 77 liver samples and ran the same differential expression and ordinal regression analyses as described for the sRNAs (above). This resulted in a limited number of significantly implicated genes (∼700 genes at BH p-adjust < 0.05), which could be attributed to reduced RIN, and for the following analysis, a higher significant threshold (BH p-adjust < 0.1) was used. At this threshold, 1256 significant genes were identified across the five analyses (Fig. 7A, Table S6). Overlap analysis of DE genes showed a 72% similarity between differential expression and ordinal regression results associated with fibrosis, and all but one significant DE genes have consistent directionality across comparisons (Fig. 7B). To validate our results, we compared fold-changes to those of another independent dataset (9) and observed strong correlations for both MASH vs MASL and advanced fibrosis (respectively r = 0.33, p-value < 10^-10^, and r = 0.44, p-value < 10^-10^, Fig. 7C-D). This independent study reported 19 and 18 genes that are predictive for NAS and fibrosis scores, most of which are similarly regulated in our dataset (red dots on Fig. 7C-D). Six of them had a significant correlation with fibrosis (labeled on Fig. 7D). Gene enrichment analyses were performed using GO-Biological Processes, KEGG and Reactome databases and many disease-relevant pathways were differentially expressed (Fig. S7, Table S7). These included repression of metabolic pathways with high fibrosis such as amino acid catabolism (p-adjust = 2×10^-19^), insulin resistance (p-adjust = 0.02), glycolysis (p-adjust = 7×10^-7^), and fatty acid metabolism (p-adjust = 0.01). Importantly, this last metabolic pathway, as well as the cholesterol metabolic pathway and PPAR pathway are inverted and showed an up-regulation associated with high NAS (p-adjust respectively 1×10^-3^, 3×10^-4^, and 0.02). We also found expected pathways associated with fibrosis activated in our dataset: focal adhesion (p-adjust = 5×10^-6^), response to TGF-beta (p-adjust = 2×10^-6^), and collagen fibril organization (p-adjust = 6×10^-11^) which were consistent with the results of gene enrichment analysis from our comparative external study (Fig. S7). The general overlap, in terms of DE genes and dysregulated pathways, between this study and an independent liver biopsy dataset supports the robustness and reliability of our results.

**Fig. 7.**
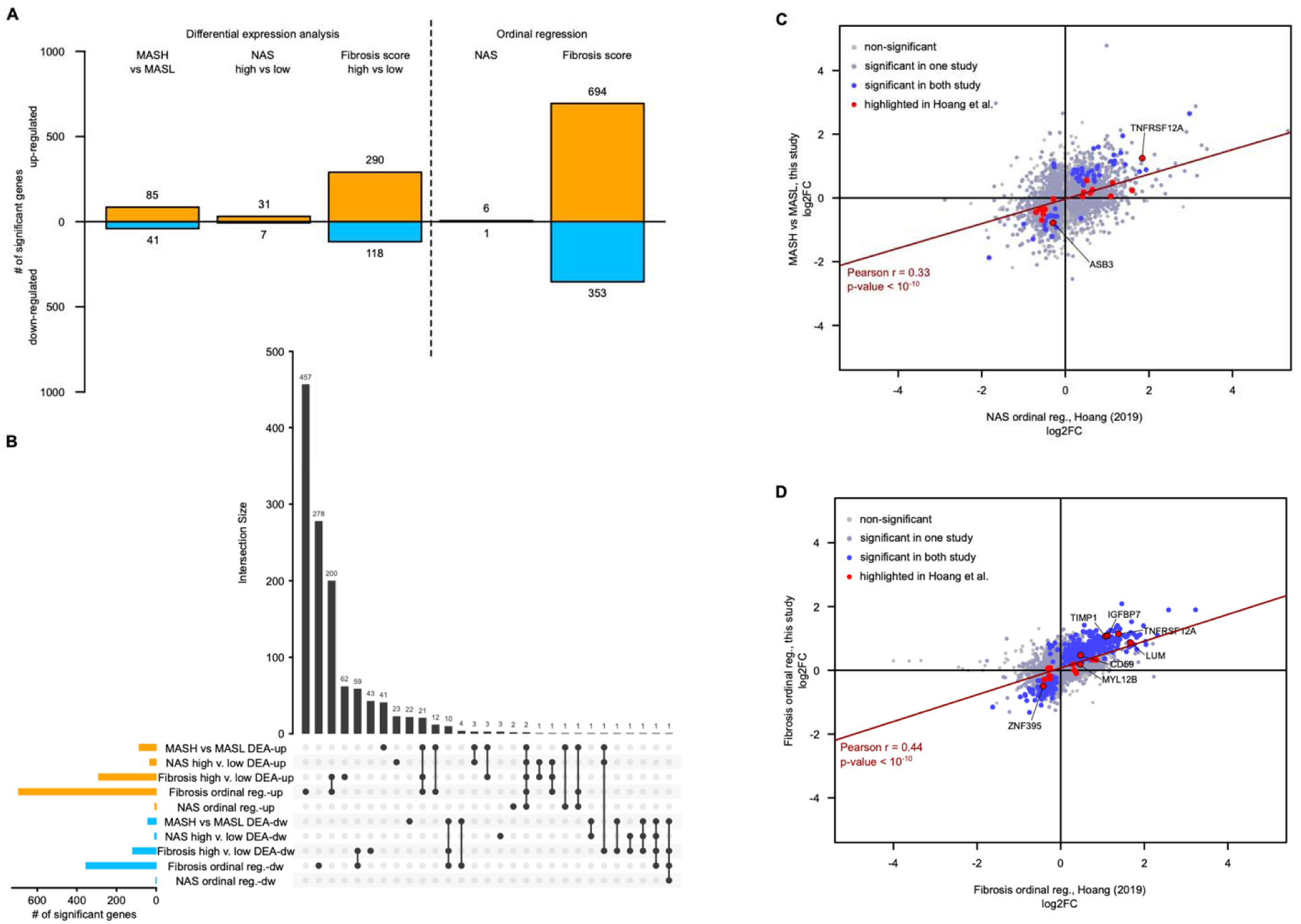
mRNA differential expression analysis. (**A**) Number of significant genes in each of the three comparator groups and two ordinal regression analyses. (**B**) Overlap analysis of significant genes across differential analyses using upset plot. (**C-D**) Inter-study comparison of significant genes associated with MASH (C), or liver fibrosis (D). Both cases showed highly significant positive correlation between the log2FoldChange across studies.

Gene expression results were used to provide potential regulatory networks for each selected sRNA and mechanism of action associating them to either MASH or Fibrosis severity. For each sRNA, the 2-8 seed sequence was used to obtain custom or canonical target lists from TargetScan (52, 53) and gene enrichment analysis was used to associate these targets to specific pathways. A total of 33 pathways, 15 for MASH and 27 for fibrosis, were selected from the dysregulated pathways presented above, as well as a set of pathways associated with liver disease or fibrosis from an in-house AI-generated knowledge graph dataset (Fig. 8A-B, Table S8). The number of targeted genes varied widely across isomiRs, from 3813 targets for the CAGUGGC miR-27b-3p seed to 10 for the AACGCCA miR-122-3p seed. Notably, differences in targeted pathways can be observed across canonical seeds and shifted seeds from 5’ NTAs or 5’ TADs within the same miRNA family. Three types of targeted genes were considered for relevant mechanisms of action: (i) differentially expressed genes involved in these 33 pathways, (ii) genes reported as strongly implicated in MASH or liver fibrosis using an AI generated knowledge graph (30 for NAS and 25 for fibrosis, Table S9), and (iii) the predictive genes from the external study (9) (19 for NAS and 18 for fibrosis, highlighted in red Fig. 7C-D). Regulatory networks were generated for all selected sRNAs (Table S10), confirming already proposed mechanisms of action, such as INSIG1 regulation by miR-363 (54), or SCD regulation by miR-27a (20). Importantly, novel regulatory mechanisms were also identified in relation to the disease, involving non-canonical seed targeting.

**Fig. 8.**
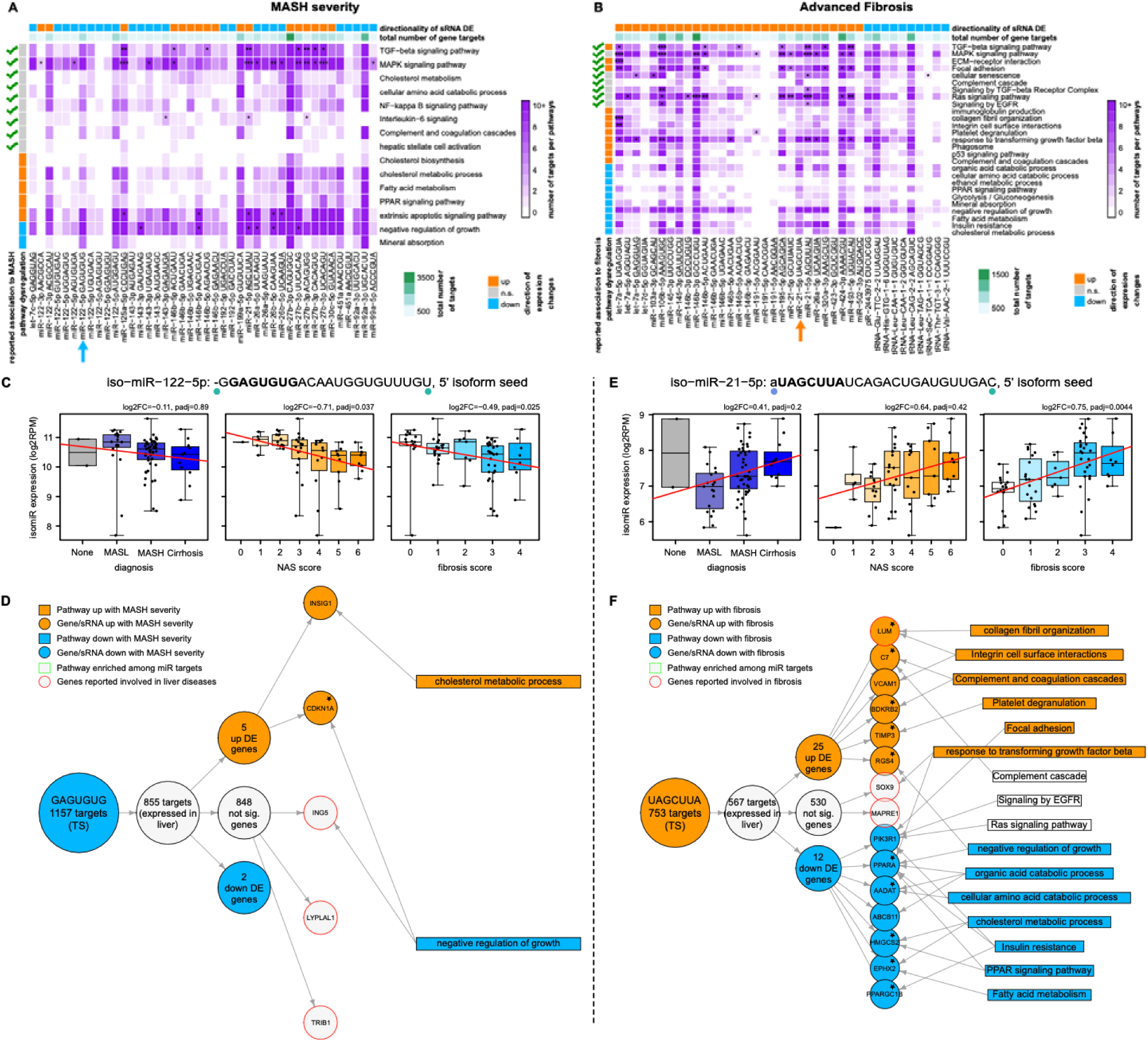
MicroRNA regulatory role in MASLD. (**A-B**) Overview of selected sRNA regulated pathways involved in MASH severity (A) and Advanced Fibrosis (B). The purple color corresponds to the number of targeted genes (based on Target-Scan) in the pathway, and stars indicates significant target enrichment (BH p-adj: <0.1: *, <0.01: **, <0.001: ***). Underlined seeds indicate canonical sequences. (**C**) Expression of the ML-selected differentially expressed isomiR from miR-122-5p family (5’ templated deletion) associated to MASH severity across diagnosis, fibrosis scores and NAS. For Fibrosis and NAS, log2FC indicates log2-FoldChange for high over low scores, and ‘padj’ indicates the BH-adjusted p-value of the ordinal regression method. (**D**) AI-informed putative regulatory network involved in MASH severity of the same miR-122-5p isomiR using Target Scan (TS) as target predictor. Stars indicate genes that are also significantly dysregulated in the same direction in the Hoang et al. study (BH p-adjusted < 0.1). (**E-F**) Same as (A-B) for an isomiR of miR-21-5p (5’ non-templated A addition) associated to fibrosis.

An isomiR from miR-122-5p family with a single 5’ nucleotide deletion was negatively correlated to NAS score (Fig. 8C), and its seed sequence had 1157 predicted targets including INSIG1 and CDKN1A, genes up-regulated with MASH severity, the former being involved in cholesterol metabolic process (Fig. 8D). Additionally, other targets such as ING5, LYPLAL1 and TRIB1, while not differentially expressed in our dataset, have been reported to be associated with liver disease (9, 55, 56). This contrasts with the canonical miR-122-5p predicted targets, consisting of 52 genes, for which we were unable to demonstrate a relationship with disease activity (Fig. S8A-B).

Similarly, the miR-21-5p isomiR carrying a 5’ adenosine addition is activated with fibrosis (Fig. 8E) and had 12 of its 753 predicted targets downregulated. Those include PIK3R1 involved in Focal Adhesion and Insulin Resistance, PPARA involved in PPAR signaling pathway and TGF-beta response, and HMGCS2 involved in cholesterol metabolism (Fig. 8F). Importantly, PPARA and HMGCS2, which have been suggested as therapeutic targets for fatty liver disease (57, 58), are not predicted targets of the canonical miR-21-3p seed (Fig. S8C-D).

Other isomiRs from miRNA families not previously reported as associated with liver disease or fibrosis, showed potentially disease-relevant regulations, especially around lipid metabolism. This is the case for an isomiR of miR-26b-5p repressed in patients with high NAS score. It targets the mRNA of ACSL4, who encode a long-chain fatty-acid-coenzyme A ligase that is activated with MASH severity (Fig. S9A-B). Another novel regulatory mechanism involves a miR-320a-2p isomiR that is positively correlated with fibrosis score. It targets RXRA, a hormone receptor required for PPARA transcriptional activity of fatty acid oxidation genes (59), which have anti-fibrotic activity in liver (60) (Fig. S9C-D).

For most of the targeted genes presented here, their expression pattern in MASLD of varying severity was confirmed in the independent dataset (e.g., CDKN1A, PPARA, HMCGS2, ACSL4, RXRA, indicated with stars on Fig. 8D-F, Fig. S8-9). These networks provide novel regulatory mechanisms associated with liver disease and cirrhosis progression and underline the importance of considering post-transcriptional modifications in miRNA function and therapeutic potential.

## DISCUSSION

The current study confirms that miRNA are the principal small RNA species in liver tissue across the histological spectrum of MASLD and that 67% of miRNAs in the liver are non-canonical isomiRs. It also demonstrates that the amount and type of post-translational modification varies across miRNA families and patients. Further, several targets in relation to disease severity, including several previously unknown targets, were identified involving non-canonical seed targeting. These novel data have several implications for the field.

First, the data provide insights on the biological complexity of MASLD and the contribution of isomiRs to the heterogeneity of the disorder. A novel finding is that miRNA isoforms constitute a substantial subpopulation within the overall miRNAs and are enriched within a relatively limited number of miRNA families. Importantly, only a small proportion of isomiRs involved alterations in the 5’ region needed for mRNA targeting indicating their potential to contribute to differential repression of genes that could impact adaptive or maladaptive responses to the upstream lipotoxic injury in MASLD and modify disease progression. The 3’ modifications within differentially expressed isomiRs were more commonly seen and, based on their known functions (27, 44–48), would be expected to impact their stability and localization.

Characterizing isomiR profiles and linking them to their mRNA targets also provide insights on both the heterogeneity of the disease and novel biology. While many differentially expressed gene targets of the isomiRs have been previously implicated in the pathobiology of MASLD e.g. PPARA and PI3KR1 involved in focal adhesion and fibrosis (18, 57, 61), several novel targets were also identified, including CDKN1A, ACSL4, and RXRA. MicroRNAs often target multiple genes creating the potential for off-target effects with miRNA-based therapeutics. Identification of isomiRs that have more specific effects on mRNA targets as noted with targets linked to lipid metabolism provides a potential explanation for the heterogeneity in gene expression which may be biologically relevant and be leveraged to attack specific targets while reducing the potential for off-target effects.

A substantially greater number of isomiRs were related to fibrosis severity than disease activity attesting to the complexity of factors regulating fibrosis progression. It is also interesting to note that while miR-122-5p family related isomiRs decreased with disease progression, others such as miR-143-3p and miR-21-5p were increased. Circulating miR-122 tends to increase with declining hepatic levels with fibrosis progression in MASLD suggesting that these are exported or released from the liver with injury and disease progression (62). Interestingly, a single isomiR of miR-122-5p was increased in those with more advanced fibrosis whereas the rest were decreased. This isomiR had modifications at its 3’ end potentially changing its cellular localization and stability.

The classic histological pattern of MASH is a predominantly zone III distribution of steatosis, hepatocellular ballooning and lobular inflammation (31). The underlying metabolic perturbations, inflammatory responses and fibrogenesis are all active energy-requiring processes. This perivenular zone also has a low oxygen tension even under normal circumstances and the increased energy requirements in MASLD may further induce tissue hypoxemia and modulate the injury and fibrogenic response. While the oxygen tension of the region has not been studied in humans with MASH, disease severity has been associated with sleep apnea which causes hypoxemia. The miR-21 family has been implicated in hypoxemia and also oncogenesis (63, 64). In the current study, a strong relationship of miR-21-5p and fibrosis severity was also noted. This raises the possibility for miR-21 to be one potential link relating the greater incidence of hepatocellular cancer in MASH with advanced fibrosis compared to early-stage disease and provides direction for future research.

Several circulating miRNAs have been evaluated as biomarkers reflective of disease severity in MASLD such as miR-34a, miR-21 and miR-122 (65). The identification of isomiRs related to varying stages of disease in the liver further opens a new avenue of research namely the identification of these isomiRs in circulation to provide a more specific way to define the severity of fibrosis in affected individuals.

## Author contributions

CB, SAH, DJH, NCF, DWS, and AJS contributed to the conceptualization of the study. The methodology was developed by CB, SAH, MRL, MAS, NCF, and DWS. The investigation was carried out by CB, GW, FM, JA, MRL, ZZ, BS, MSS, and AA. Data visualization was handled by CB, SAH, and MRL. NCF, DWS, and AJS were responsible for funding acquisition, project administration, and supervision. The original draft of the manuscript was written by CB and AJS, with subsequent review and editing by SAH, GW, MRL, NCF, and DWS.

## Supplementary data

Supplementary figure file: Fig. S1 to S9
Supplementary table file: Table S1 to S10

## Data availability

Most relevant data are available in the main text or the supplementary materials. Additional data are available upon request.

## Abbreviations

AUROC: receiver operator characteristic area under the curve
isomiRs: microRNA isoforms
LLMs: Large Language Models
MASLD: metabolic dysfunction-associated steatotic liver disease
MASL: metabolic dysfunction associated steatotic liver
MASH: metabolic dysfunction associated steatohepatitis
miRNA: microRNA
NAS: metabolic dysfunction-associated steatotic liver disease (previously NAFLD) activity score
NTA: Non-Templated Addition
PPAR: peroxisome proliferator-activated receptor TAD: Templated Addition or Deletion

