## Supplemental Figures for "Hepatic isomiR landscaping reveals new biological insights into metabolic dysfunction in steatotic liver disease"

### Supplemental Figures S1-S9

### Hepatic isomiR landscaping reveals new biological insights into metabolic dysfunction in steatotic liver disease

Christian Brion[1], Stephen A. Hoang[2], Guangliang Wang[1], Faridodin Mirshahi[2], Jessie Ang[1], Matthew R. Long[1], Zheng Zhu[1], Bhanu Sakhamuri[1], Molly A. Srour[1], Mohammad S. Siddiqui[2], Amon Asgharpour[2], David J. Hayes[1], Neal C. Foster[1], David W. Salzman[1*], Arun J. Sanyal[2*]

1. Gatehouse Bio Inc.; 22 Strathmore Road, Natick, MA, 01760, USA.

2. Division of Gastroenterology, Hepatology and Nutrition, Department of Internal Medicine, Virginia Commonwealth University School of Medicine; Richmond, VA, 23298, USA.

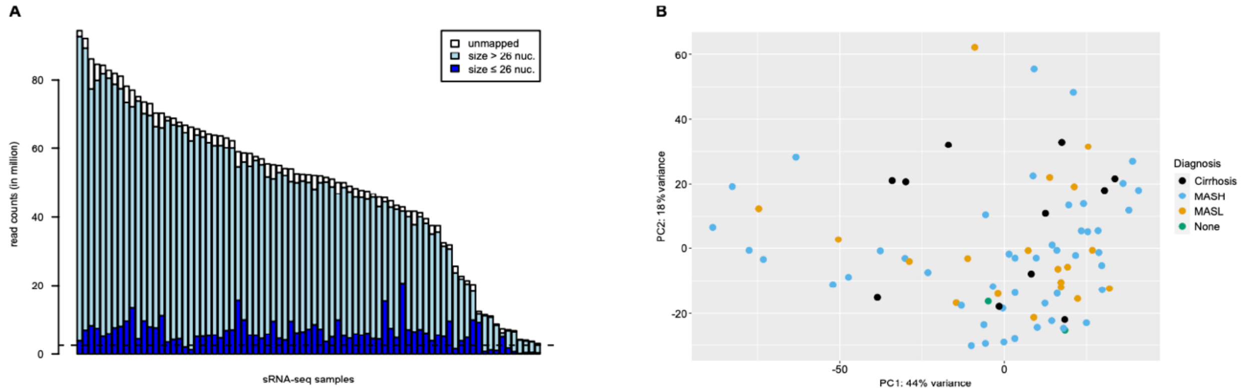


**Fig. S1. sRNA sequencing output.** (**A**) Sequencing depth and size selection for sRNA sequencing. (**B**) Principal Component Analysis of the sRNA sequencing data using the 500 most variable features. No clear cluster and outliers were identified.

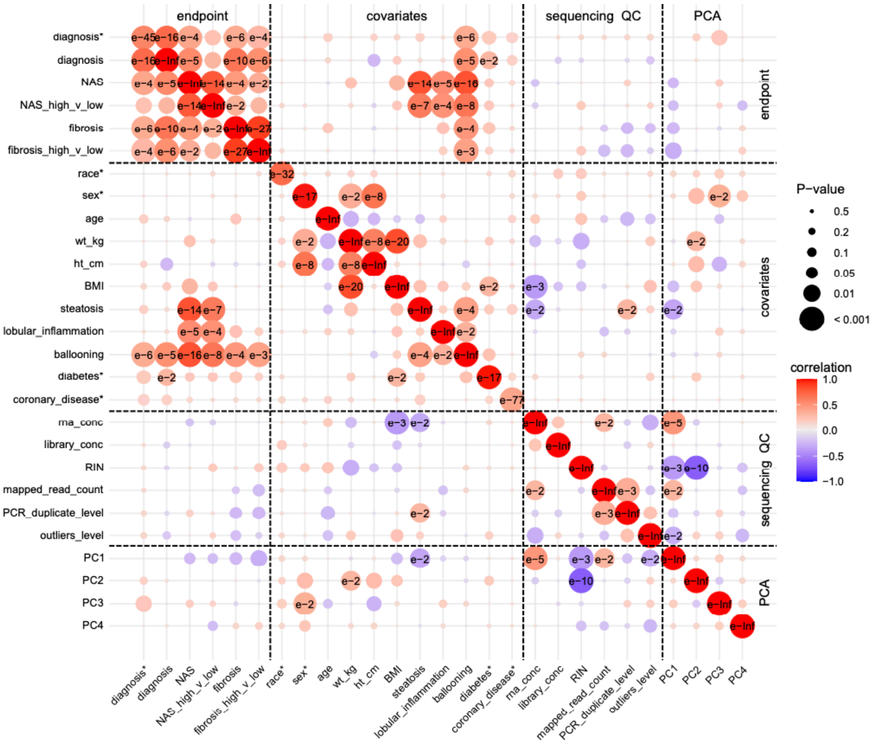


**Fig. S2. Pairwise meta data covariate comparison.** Factor variables are indicated with a star. Continuous to continuous, factor to continuous, and factor to factor correlations values were calculated by Spearman's correlation method, Kruskal-Wallis based Cramer's V, and Pearson's Chi2 based Cramer's V, respectively.

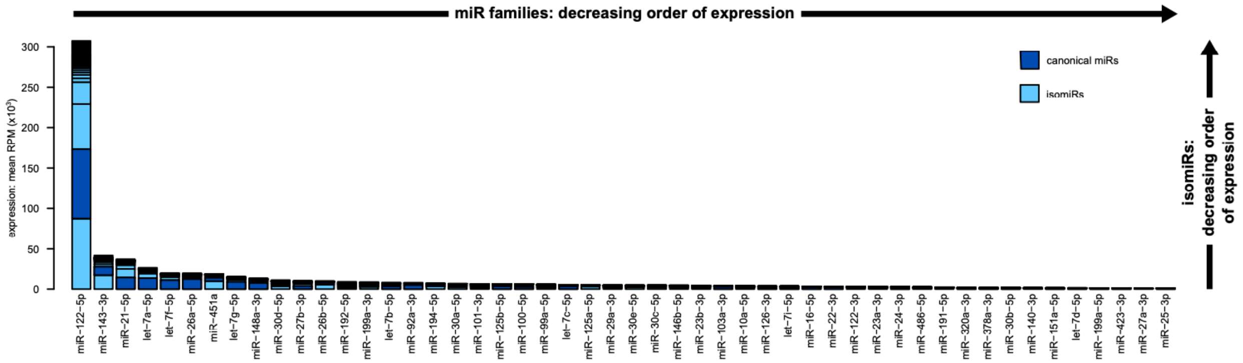


**Fig. S3.** Distribution of the expression of the 50 most highly expressed miRNA families.

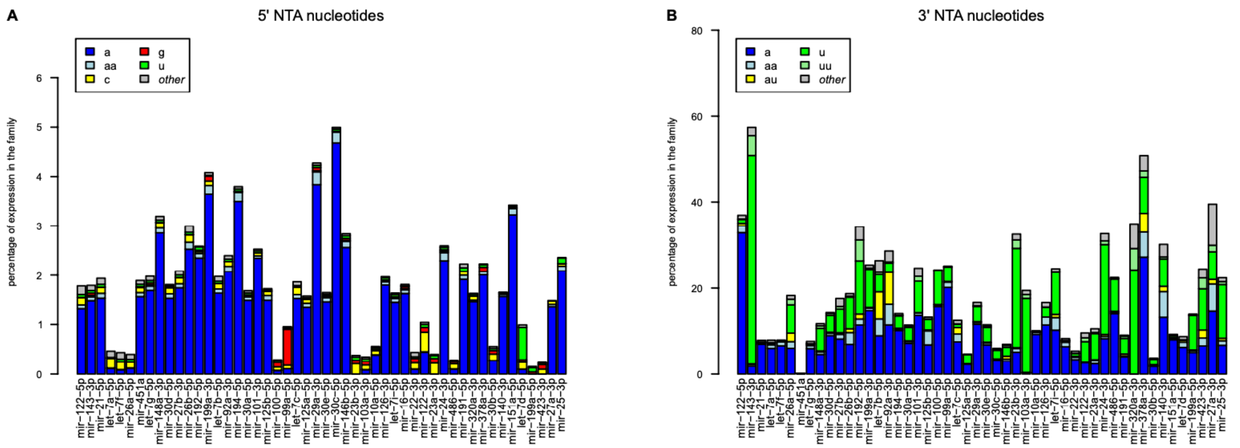


**Fig. S4.** Type and occurrence of non-templated additions (NTA) at the 5’ (A) or 3’ (B) ends of the isomiRs for each of the 50 most highly expressed miRNA families.

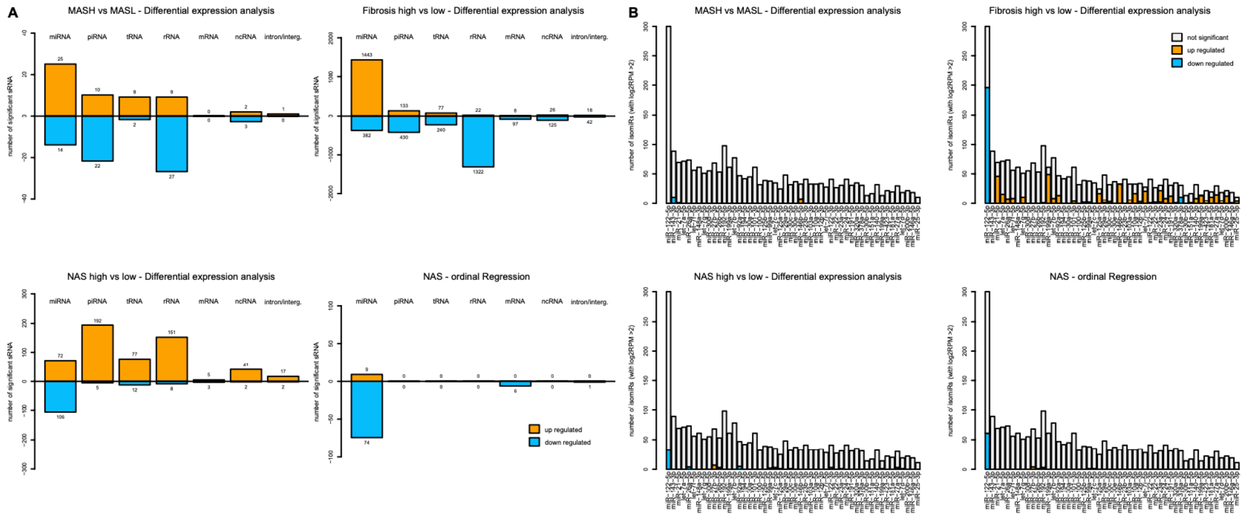


**Fig. S5.** (**A**) Breakdown of the number of significant sRNA by type across four comparative methods (MASH vs MASL DEA, NAS high vs low DEA, Fibrosis high vs low DEA, and NAS ordinal regression analysis). (**B**) Number of significant isomiRs within each 50 most highly expressed miRNA families across the same four comparative methods.

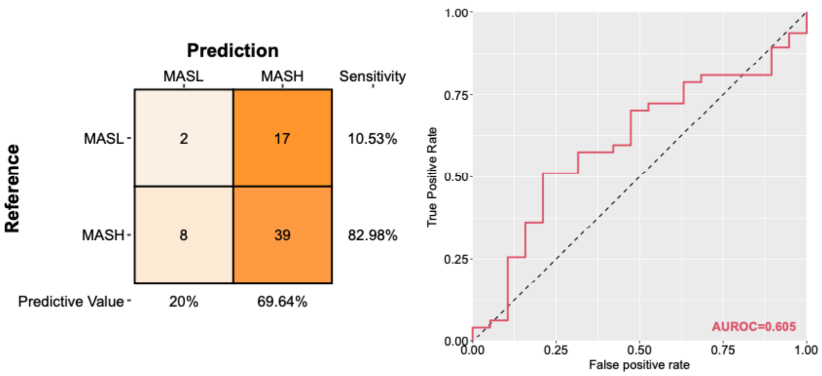


**Fig. S6.** Machine learning predictive outcome for MASH vs MASL with confusion matrix (left) and ROC curve (right).

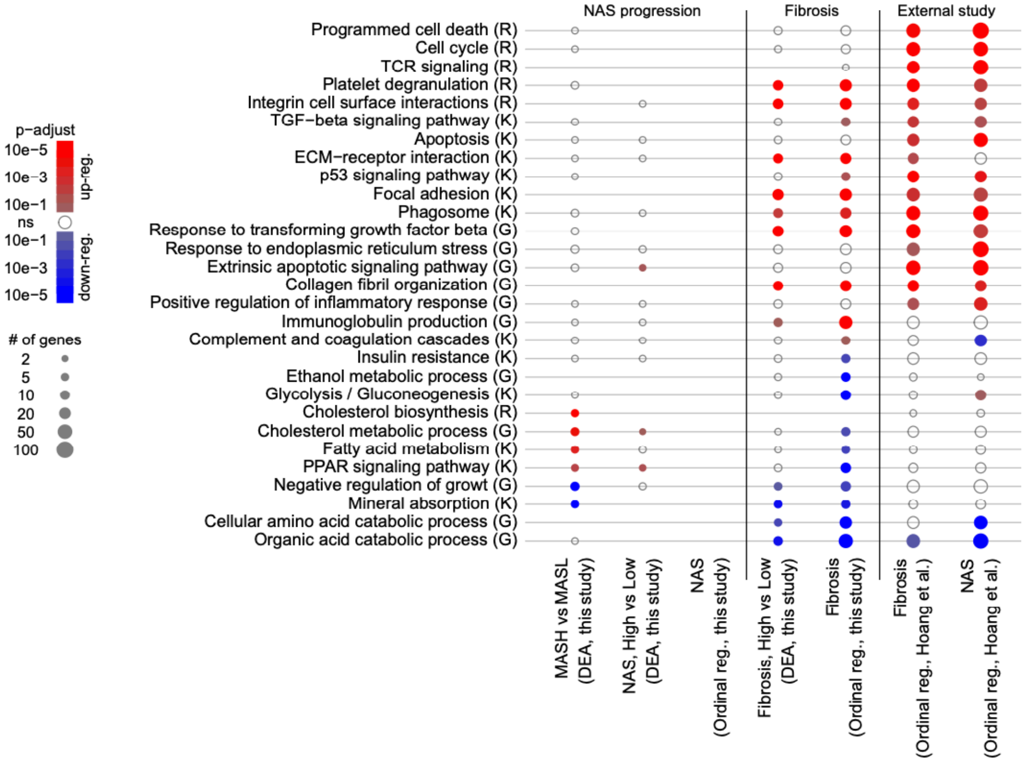


**Fig. S7. Inter-study comparison of gene set enrichment analysis.** Significantly differentially expressed genes across each comparative methods were considered with the following threshold: BH p-adjust < 0.1 from our study, and < 0.05 from Hoang et al. study. We used KEGG (K), Reactome (R), and GO: Biological Process (G) repositories. NAS ordinal regression did not generate enough significant genes for successful enrichment analysis.

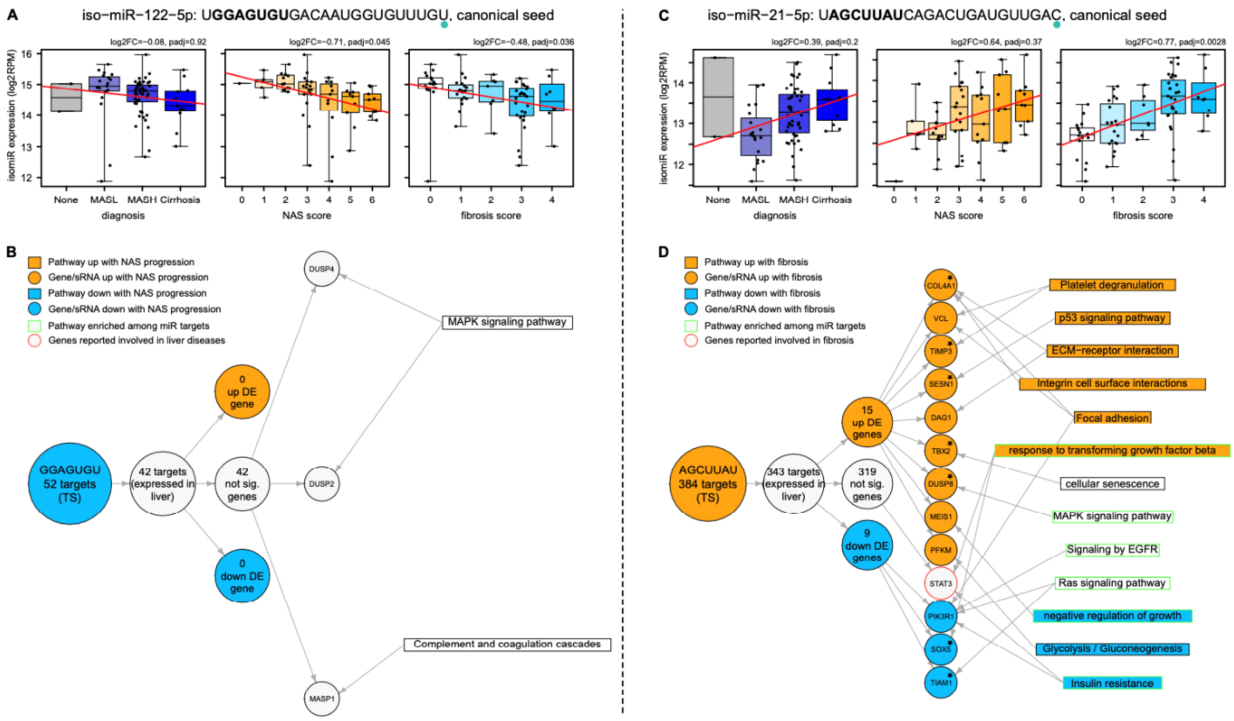


**Fig. S8. MicroRNA regulatory role in MASLD.** (**A**) Expression of a ML-selected differentially expressed isomiR from miR-122-5p family (canonical seed) associated with MASH severity across diagnosis, fibrosis scores and NAS. For Fibrosis and NAS, log2FC indicates log2-FoldChange for high over low scores, and ‘padj’ indicates the BH-adjusted p-value of the ordinal regression method. (**B**) AI-informed putative regulatory network involved in MASH severity of the same miR-122-5p isomiR using Target Scan (TS) as target predictor. Stars indicate genes that are also significantly dysregulated in the same direction in the Hoang et al. study (BH p-adjusted < 0.1). (**C-D**) Same as (A-B) for an isomiR of miR-21-5p (canonical seed) associated to advanced fibrosis.

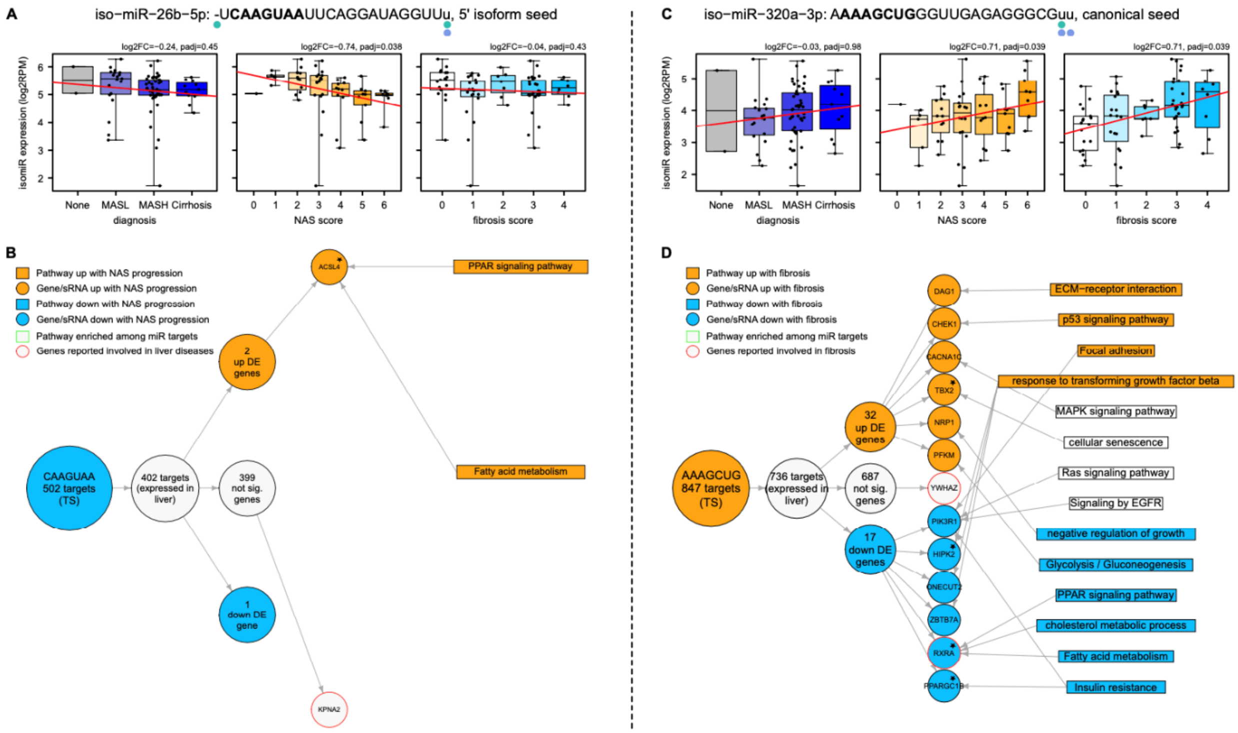


**Fig. S9. MicroRNA regulatory role in MASLD.** (**A**) Expression of a ML-selected differentially expressed isomiR from miR-26b-5p family (one 5' deletion) associated with MASH severity from across diagnosis, fibrosis scores and NAS. For Fibrosis and NAS, log2FC indicates log2-FoldChange for high over low scores, and ‘padj’ indicates the BH-adjusted p-value of the ordinal regression method. (**B**) AI-informed putative regulatory network involved in MASH severity of the same miR-26b-5p isomiR using Target Scan (TS) as target predictor. Stars indicate genes that are also significantly dysregulated in the same direction in the Hoang et al. study (BH p-adjusted < 0.1). (**C-D**) Same as (A-B) for an isomiR of miR-320a-5p (canonical seed) associated to advanced fibrosis.
